# Computer-aided drug screening of anti-*Neobenedenia melleni* drugs based on annexin B1 in farmed pearl grouper (*Epinephelus fuscoguttatus*♀ × *Epinephelus lanceolatus* ♂)

**DOI:** 10.64898/2026.08.23.746516

**Authors:** Longkun Gao, Wei Luo, Yanru Guo, Ying Yan, Guanhai Li, Qin Yu, Mingzhu Liu, Erlong Wang, Pengfei Li, Tianqiang Liu

## Abstract

Monogenean capsalids of the genus Neobenedenia are widespread parasites of wild and farmed marine fish and represent a major threat to grouper mariculture in China, whose production reached approximately 294,000 tonnes in 2025. The development of effective drugs to control and prevent these infections is therefore urgently needed. Annexins, which have been identified in Neobenedenia and other parasites, differ markedly from their host counterparts, making them potentially attractive targets for antiparasitic therapeutics. Here, we performed a computer-aided drug discovery screen, employing *Neobenedenia melleni* annexin B1 as the molecular target against a library of 1,456,161 small molecules. The three-dimensional structure of annexin B1 was first predicted using AlphaFold 3, SWISS-MODEL, and I-TASSER, and the most accurate model (AlphaFold) was selected for structure-based virtual screening. *In vivo* validation of eleven compounds identified abamectin (Aba) as the most effective anti-Neobenedenia agent, achieving complete parasite elimination at 0.16 mg/L. Owing to its low toxicity to the host grouper (24-h LC_50_ = 0.254 mg/L), abamectin was selected for further investigation. Abamectin exhibited potent anthelmintic activity against *N. melleni*, with a 24-h bath exposure yielding an EC_50_ of 0.033 mg/L and complete parasite elimination at 0.16 mg/L, corresponding to a therapeutic index of approximately 7.7. To elucidate the antiparasitic mechanism, we performed long-timescale (1000 ns) molecular dynamics simulations of the annexin B1-abamectin complex, enabling atomic-level analysis of the essential protein motions involved in their interaction. The interaction profile between annexin B1 and abamectin was dominated by hydrophobic contacts and water bridges, involving residues TYR-210, GLU-214, GLU-244, and SER-247, which path a way for further drug optimization.

## Introduction

Grouper (Epinephelus spp.) aquaculture is among the most valuable and rapidly expanding sectors of marine finfish culture in tropical and subtropical Asia, and China now accounts for the majority of global production, which reached approximately 294,000 tonnes in 2025.[1] The intensification of cage and pond culture has been accompanied by a parallel rise in disease outbreaks, particularly those caused by monogenean ectoparasites.[2] Among these, the capsalid monogenean *Neobenedenia melleni* is one of the most damaging, owing to its exceptionally broad host range and its capacity to proliferate rapidly in high-density culture systems.[3] *N. melleni* attaches to the skin, fins, and eyes by means of a posterior haptor, feeds on epithelial cells and blood, and induces extensive integumentary damage, osmotic stress, and secondary bacterial infections that frequently culminate in mass.[4] The production of numerous, highly resistant eggs further complicates control, because reinfection occurs rapidly once eggs hatch, necessitating repeated therapeutic intervention.[5]

The rational discovery of new control agents requires well-characterized, parasite-specific molecular targets. Annexins, a conserved family of calcium- and phospholipid-binding proteins present in virtually all organisms, have emerged as promising candidates in this regard.[6] Members of this superfamily are classified into families A-E according to their phylogenetic distribution, with family A in vertebrates and family B in protostome and deuterostome invertebrates, fungi, and other non-vertebrate.[7] Importantly, parasite annexins are frequently and markedly divergent from their host counterparts, a distinction that has been proposed to render them attractive, species-selective targets for antiparasitic drugs and vaccines.[8] Consistent with this notion, our phylogenetic analysis of all available annexin sequences of monogeneans revealed that *N. melleni* possesses annexin B1, which is clearly distinct from the annexin repertoire of its grouper hosts (Epinephelus spp.). This finding indicates that parasite annexin B1 represents a rational, target-specific candidate for structure-based drug screening against *N. melleni*.

Despite the economic importance of monogenean infections, the chemical arsenal available for their control remains limited. Current treatments rely almost exclusively on a small number of bath-administered chemicals-freshwater or hyposaline dips, formalin, hydrogen peroxide, praziquantel, and a few avermectin-class, most of which were discovered decades ago through low-throughput phenotypic screens in animal models of infection.[9] This narrow repertoire is increasingly compromised by drug resistance to broad-spectrum, raising concerns over environmental toxicity and non-target effects.[10]

Although advances in genomics and protein-structure prediction have generated an abundance of parasite sequence data, these developments have not yet been translated into new anthelmintic classes, and screening for aquatic monogeneans remains almost exclusively phenotypic and low-throughput.[11] Moreover, while parasite annexins are acknowledged in principle as attractive drug targets, no monogenean annexin has yet been validated as a druggable target through a structure-based screen.[12] Consequently, there is currently no rational, target-directed route to identify parasite-selective and environmentally benign lead compounds against *N. melleni*-precisely the combination required to circumvent resistance and host toxicity.

In the study, we performed a structure-based virtual screening campaign targeting *N. melleni* annexin B1. The three-dimensional structure of annexin B1 was first predicted independently using AlphaFold, SWISS-MODEL, and I-TASSER, and the highest-quality model (AlphaFold) was selected following rigorous validation with SAVES and MolProbity. This model served as the target for docking-based screening of a large and chemically diverse library of approximately 1.45 million compounds, using a hierarchical HTVS → SP → XP workflow combined with MM/GBSA rescoring and ADMET filtering via SwissADME and pkCSM. Eleven top-ranked, commercially available candidates were then assessed *in vivo* against naturally infected grouper. Among these, abamectin-a repurposed macrocyclic lactone-emerged as the most potent lead, achieving complete parasite elimination at 0.16 mg/L with an EC_50_ of 0.033 mg/L and a favorable therapeutic index (TI ≈ 7.7) relative to the host 24-h LC_50_ of 0.254 mg/L. Long-timescale (1000 ns) molecular dynamics simulations and MM/GBSA binding-free-energy calculations further revealed that abamectin engages annexin B1 through stable hydrophobic contacts and water-bridged polar interactions with key residues (TYR-210, GLU-214, GLU-244, and SER-247), substantiating annexin B1 as a bona fide drug target. Collectively, this work represents the structure-based screen against a monogenean parasite; it validates parasite annexin B1 as a druggable target class, and it delivers a practical, low-toxicity candidate for the control of *N. melleni* in grouper aquaculture.

## 2. Materials and methods

### 2.1. Materials Ethics statement

All animal experiments in this project were reviewed and approved by the Institutional Animal Care and Use Committee of Northwest A&F University with permission number DK20250512001.

#### Hardware and Software

Ubuntu 24.04.4 LTS operating system with 2 AMD EPYC 7R32 48-core processor, 1 Nvidia GeForce RTX4090, 16 ×32 Gb 3200 MT/s DDR4 DRAM. All needed software were listed and cited in proper position in the following experimental section.

#### Fish, parasites and materials

Groupers (*Epinephelus fuscoguttatus*♀ × *Epinephelus lanceolatus* ♂) (120±4 g) were obtained from a grouper fish farm in Fujian Province, China. 400 healthy fish (exposure to freshwater for 30 s) were temporarily cultured in a 10 m × 8 m × 1.2 m concrete pond for two weeks with a temperature of 28 ± 0.5°C in filtered natural seawater. Fish were fed with commercial fish pellet feed twice a day, amounting to 1% of their body mass. After acclimatization, fish were randomly divided into 4 groups, each with 90 fish with three repeats. All chemicals were purchased from Macklin Technology Co. Ltd., and used without further purification. A new stock solution of each test chemicals was fresh-prepared for each test. Collection of eggs of *Neobenedenia melleni* from naturally infected fish, methods for eggs hatchery and establishment of *Neobenedenia melleni* infected groupers were described by Ellis et. al.,[13, 14]. Fish were treated with test chemicals when average infection level reached ≥ 5 monogeneans per fish.

### 2.2. Sequences collection and phylogenetic analysis

Annexin amino acid sequences of different grouper species, including *Epinephelus coioide*, *Epinephelus bruneus*, *Epinephelus fuscoguttatus*, *Epinephelus moara*, *Epinephelus lanceolatus*, *Neobenedenia melleni*, and other Monogeneans (*Cichlidogyrus casuarinus*, *Microcotyle sebastis*) were downloaded from NCBI[15] as of May, 15^th^ 2026. The phylogenetic analysis for *annexin* was inferred from the amino acid sequence alignments by using MEGA12.[16] The best-fit amino acid substitution model was selected automatically by ModelFinder based on the Bayesian information criterion (BIC), which identified WAG+G as the optimal model. Branch supports were assessed with 1000 ultrafast bootstrap (UFBoot) replicates. The resulting tree was visualized and annotated in iTOL.[17]

### 2.3. Structure-based virtual screening of drug datasets

Ligands preparation: For annexin B1-based virtual screening, drug libraries from ZINC natural compounds dataset, TargetMol (L6020 dataset), Maybridge drug-like diversity library, Enamine calcium ion channel library, and Specs academic compound library of more than 1.4 million molecules, were used for screening to find potential anti-parasite compounds. Openbabel software[18] was applied to prepare the compounds, which could use for the following molecular docking.

Proteins preparation: 3D structural model for annexin of *Neobenedenia melleni* were done using AlphaFold Server[19], SWISS-MODEL[20], and I-TESSAR[21] in their default parameters. Structural quality assessment of the predicted models of annexin was analyzed by RPOCHECK, ERRAT, WHATCHECK, and VERIFY in SAVES server (https://saves.mbi.ucla.edu/), and MOLprobity.[22] The best-mapped structure with the least number of residues in unfavored region was chose and used for the following virtual screening to identify compounds that bind to annexin B1.

Drug screening process: After the abovementioned procedures, protein preparation wizard were used to prepare protein macromolecule, including removing Het atoms and all water molecules, addition of polar hydrogen, and examination of any missing residues. Finally, Kollman’s charges were applied to neutralize protein, and Gasteiger charges were calculated. For virtual screening, a 1 Å of grid box was generated by sitemap and receptor grid generation, centering at the point of calcium binding site. The central xyz axis of the grid box was set to 100 Å × 100 Å × 100 Å. The virtual screen process was repeated twice to ensure the accuracy of molecular docking results, with the exhaustiveness setting to 5, and the number of nodes setting to 20. Other parameters were set as default and the result obtained was analyzed manually by PyMol[23] and LigPlot+.[24] A RMSD value of less than 2 Å indicates the reliability of the docking pose.

Following virtual screening, molecules with lower docking score < −7.5 kcal/mol were subsequently evaluated with deep learning models to predict drug-target affinity and to assess the stability of the corresponding protein-ligand complexes. A total of 11 commercially available top hits from each group was selected for the next in vivo anti-parasite validation.

SwissADME-based pharmacokinetic evaluation: The pharmacological potential and ADME (absorption, distribution, metabolism, and excretion) properties of the selected compounds were evaluated as part of the drug development workflow.[25] Their ADME profiles were obtained using the SwissADME server (http://www.swissadme.ch/), while the pkCSM database (https://biosig.lab.uq.edu.au/) was employed to predict compound toxicity.[26]

### 2.4. Acute toxicity test

To determine safe immersion concentrations for top hit compounds, acute toxicity experiments were performed using groupers as host fish. The tests were carried out in 300-L glass tanks containing 260 L aerated natural seawater maintained at 28.0±0.5 °C. Each tank was stocked with 10 groupers, and different concentrations of the compound stock solutions were added to the water to predetermined concentration. A control group was exposed to 0.2% DMSO. Individuals showing stress signs or abnormal, erratic swimming behavior were immediately removed from the tanks. Mortality was recorded after 48 h of continuous exposure, and each treatment was conducted in triplicate.

### 2.5. In vivo anthelmintic activity

Six groupers were randomly selected from the infection pond and transferred into separate experimental tanks measuring 140 × 45 × 40 cm, each filled with 200 L of the test solution and maintained at 28.0 ± 0.5°C for 24 h. A negative control group was treated with 0.2% DMSO to account for any solvent-related effects on the parasites. Parasite numbers on the body surfaces of each fish were counted under a stereomicroscope at 0 h (before treatment) and 24 h (after treatment). Anthelmintic efficacy (E) for each compound was then calculated according to the method described in the article[27], using the formula E = (N − NS)/N × 100% when N > NS, and E = 0 when N ≤ NS, where N represents the initial number of *N. melleni* on the body surfaces before treatment and NS denotes the number of surviving parasites after treatment.

### 2.6. Molecular dynamics (MD) simulation and post-MD MM-GBSA

Molecular dynamics simulations were performed using the Desmond platform[28] for the top-ranked ligands selected based on their ADMET profiles. Each ligand-annexin B1 complex was solvated in an orthorhombic box of dimensions 10 × 10 × 10 Å containing TIP3P water molecules. The system was neutralized and adjusted to a physiological salt concentration of 150 mM by adding Na^+^ ions and Cl^−^ ions. Energy minimization was then carried out using the OPLS4 force field[29], followed by equilibration under NVT and NPT ensembles with a total of 41928 atoms. The production run was extended to 1000 ns at a constant temperature of 300 K and a pressure of 1.01325 bar, with coordinates saved every 1000 ps. Long-range electrostatic interactions were computed using the Particle Mesh Ewald method, and a cutoff radius of 9.0 Å was applied to Coulomb interactions. Water molecules were represented by the simple point charge model. A total of 1,000 frames were collected during the simulation and subsequently analyzed with the Desmond Simulation Interaction Diagram tool. To analyze the simulation trajectories, the dynamic protein-ligand interaction diagrams, root-mean-square deviation (RMSD), and root-mean-square fluctuation (RMSF) were calculated, and each trajectory was also inspected visually. To further validate the MD results, the binding free energies (ΔG_bind_) of the complex was computed over the entire simulation trajectories using the MM/GBSA method.[30]

### 2.7. Statistics

Statistical analyses were performed using Origin2024 software (Origin Lab Corp, USA). Significance was determined by analyzing the data using one-way ANOVA followed by a *post hoc* Tukey test. All data were presented as mean ± SD. LC_50_ was calculated as growth/sigmoidal with DoseResp Function.

## Results and Discussion

### Phylogenetic analysis of annexin from host fish and Monogeneans

The annexin protein superfamily was first identified about almost half a century ago. Human annexin A7, originally referred to as synexin, was the earliest member to be isolated and purified,[31] while human annexins A1 and A2, previously named lipocortin and calpactin respectively, were the first to be successfully cloned.[32, 33] The term “annexin” was formally introduced in 1990 to designate this growing superfamily. Since then, the 12 annexins shared by vertebrates have been grouped into the annexin A family and assigned the nomenclature annexins A1-A13 (or ANXA1-ANXA13), with A12 deliberately omitted from the official naming system. In non-vertebrate organisms, annexins are divided into several additional families: family B occurs in invertebrates, family C in fungi and certain unicellular eukaryotes, family D in plants, and family E in protists. Moreover, more than 40 other annexin subfamilies have been identified and still await formal taxonomic assignment.[34] The annexin B family, which occurs in both protostome and deuterostome invertebrates, has experienced numerous lineage-specific duplication events.[35] As a result, more than 20 distinct subfamilies have arisen, and these differ across clades-as well as from vertebrate annexins-in terms of gene architecture, protein structure, and chromosomal organization.[36] These findings also were validated by the phylogenetic analysis of all available annexin sequences in NCBI for the genus of *Epinephelus* and all monogeneans. As shown in Figure 1 of the maximum likelihood tree, *N. melleni* only has annexin B1, while other parasites either has annexin B or annexin A. They all shared a distinct difference from that of Epinephelus sp. The findings theoretically indicate that annexin B1 in *N. melleni* could be a potential drug target candidate for structure-based drug screening against *N. melleni*.

**Figure 1.**
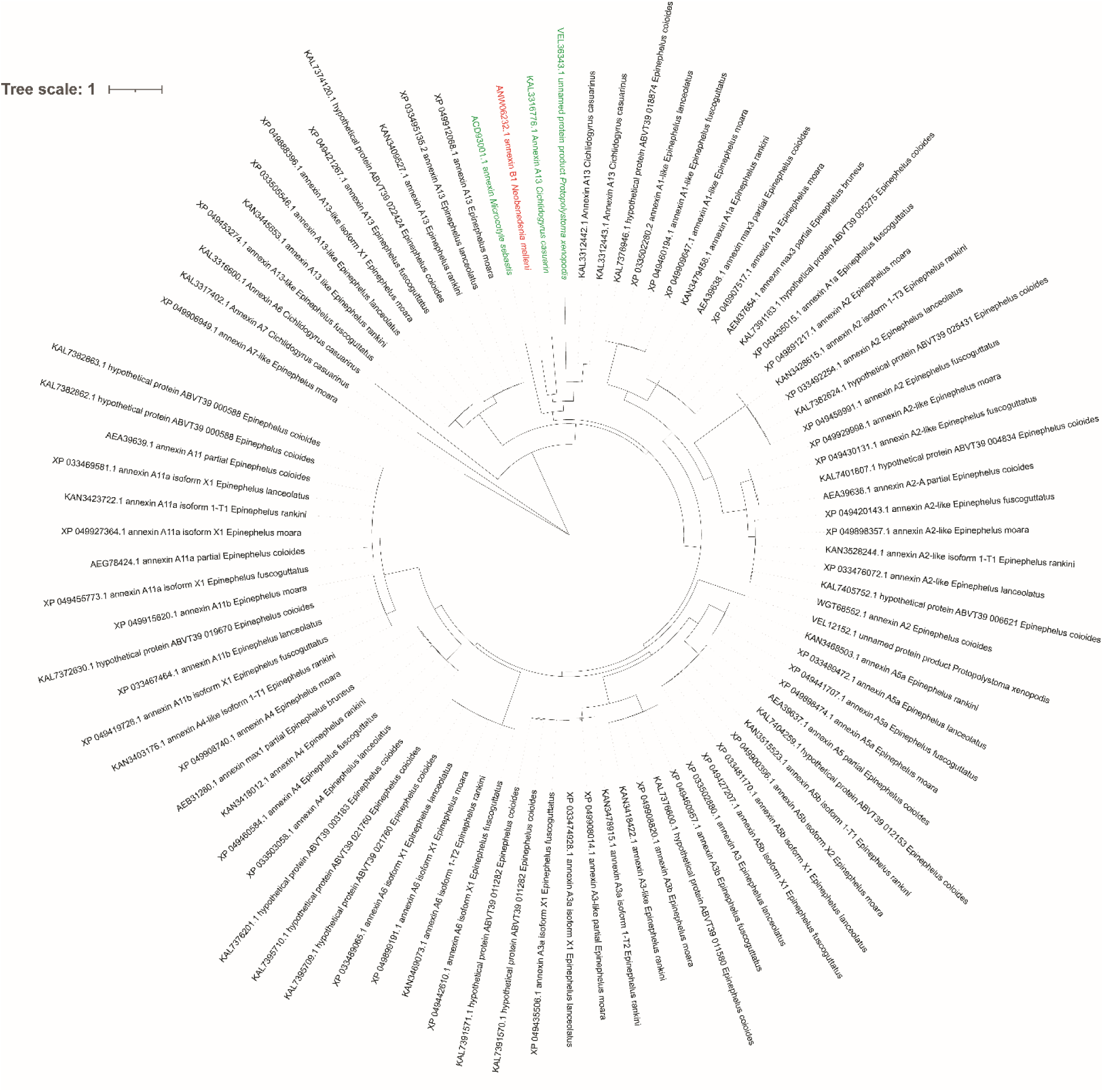
Phylogenetic tree of different type of annexin in the genus of *Epinephelus* and monogeneans from all available protein sequences from NCBI database. Names and Gene ID of the annexins are shown, with *N. melleni* labeled in red, and all other monogeneans (*C. casuarinus, M. sebastis, P. xenopodis*) labeled in green.

### Results of virtual screening based on the annexin B1 of *N. melleni*

For the target protein, the amino acid sequence of ANX B1was uploaded to the online server of AlphaFold, Swiss-MODEL, and I-TASSER, respectively. Structural models were predicted on their default parameters. Top ranked pdb file of ANX B1 from the above-mentioned three protein structure prediction tools was chosen for comprehensive quality assessments based on SAVES (https://saves.mbi.ucla.edu/) and Mol-Probity (https://molprobity.biochem.duke.edu/index.php) to rigorously evaluate the stereochemical reliability of each predicted structure. Main parameters that could reflect the quality was summarized in Table 1, which showed that ANX B1 predicted by both AlphaFold and SWISS-MODEL pass by the VERIFY analysis, while that of I-TASSER fail. This phenomenon is typically observed in multi-template threading approaches when the target sequence shares low sequence identity with the available structural templates, leading to an incorrect global fold or misplacement of secondary structural elements.[37, 38] Unlike AlphaFold, which incorporates evolutionary covariance and attention mechanisms to capture long-range residue interactions, I-TASSER relies heavily on fragment assembly guided by threading alignments.[39] Consequently, ambiguous alignment regions are prone to generating non-native side-chain orientations and incorrect backbone topology. The resulting model, while possessing a global score that passes certain criteria, fails to meet the stringent sequence-environment compatibility checks implemented by VERIFY3D, ultimately rendering it structurally implausible for reliable downstream docking studies. For ERRAT analysis, structures predicted by AlphaFold and SWISS-MODEL yielded excellent scores of 99.71 and 100.00, respectively, whereas the I-TASSER model obtained a remarkably low score of 13.49, indicating poor atomic packing and a high probability of steric clashes. Crucially, MolProbity analysis highlighted the superior dihedral angle distribution of the AlphaFold2 model. As shown in Figure 2, it achieved a Ramachandran favored score of 98.27% and an extremely favorable rotamer distribution (99.66%). Overall, structure predicted by AlphaFold demonstrated the most robust backbone geometry, displaying only minimal deviations, thus was chosen for the following structure-based drug screening.

**Figure 2.**
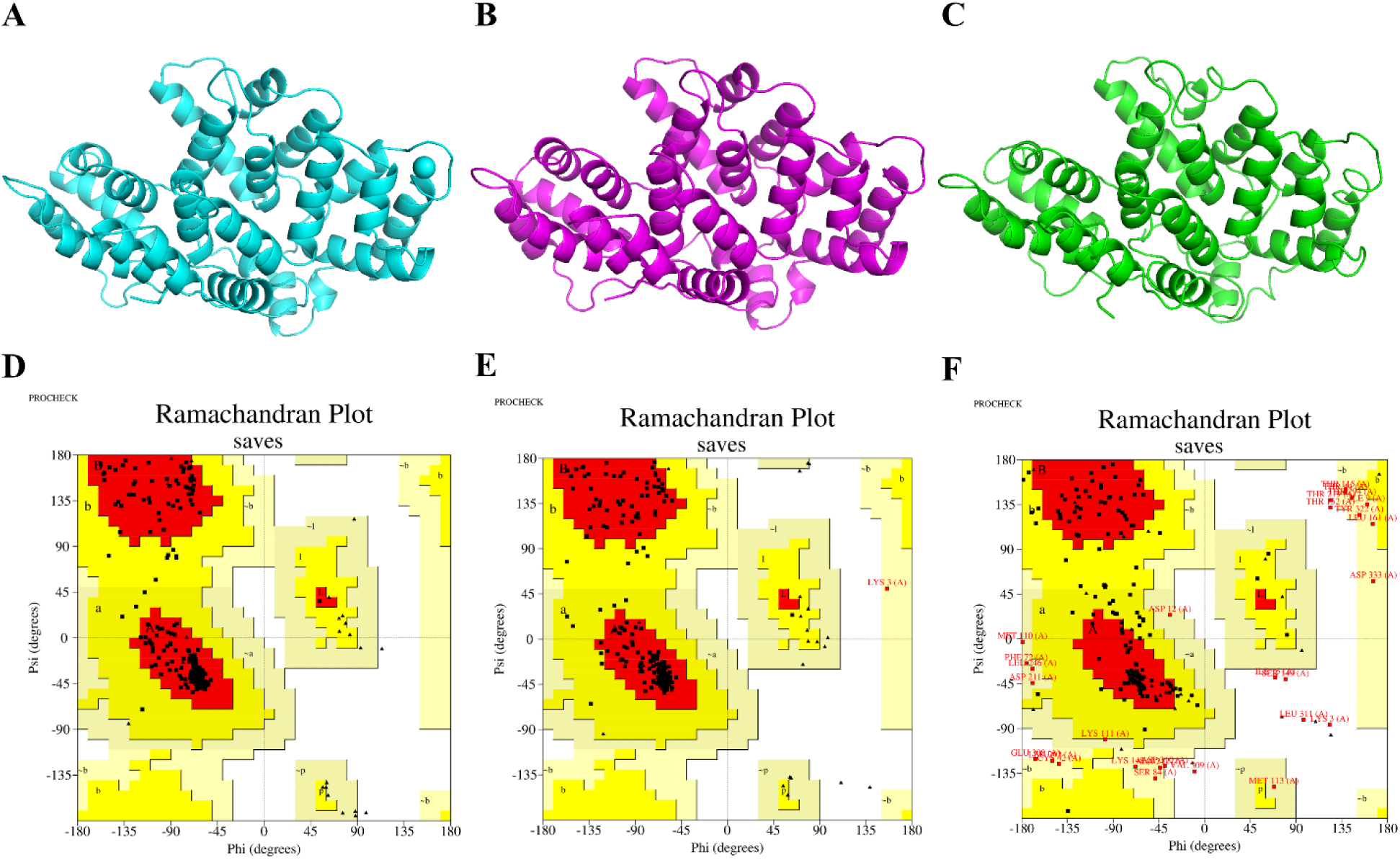
The 3-D structures of ANX B1 predicted by AlphaFold (A), Swiss-MODEL (B), I-TASSER (C). The corresponding Ramachandran plot of AlphaFold (D), Swiss-MODEL (E), I-TASSER (F).

**Figure 3.**
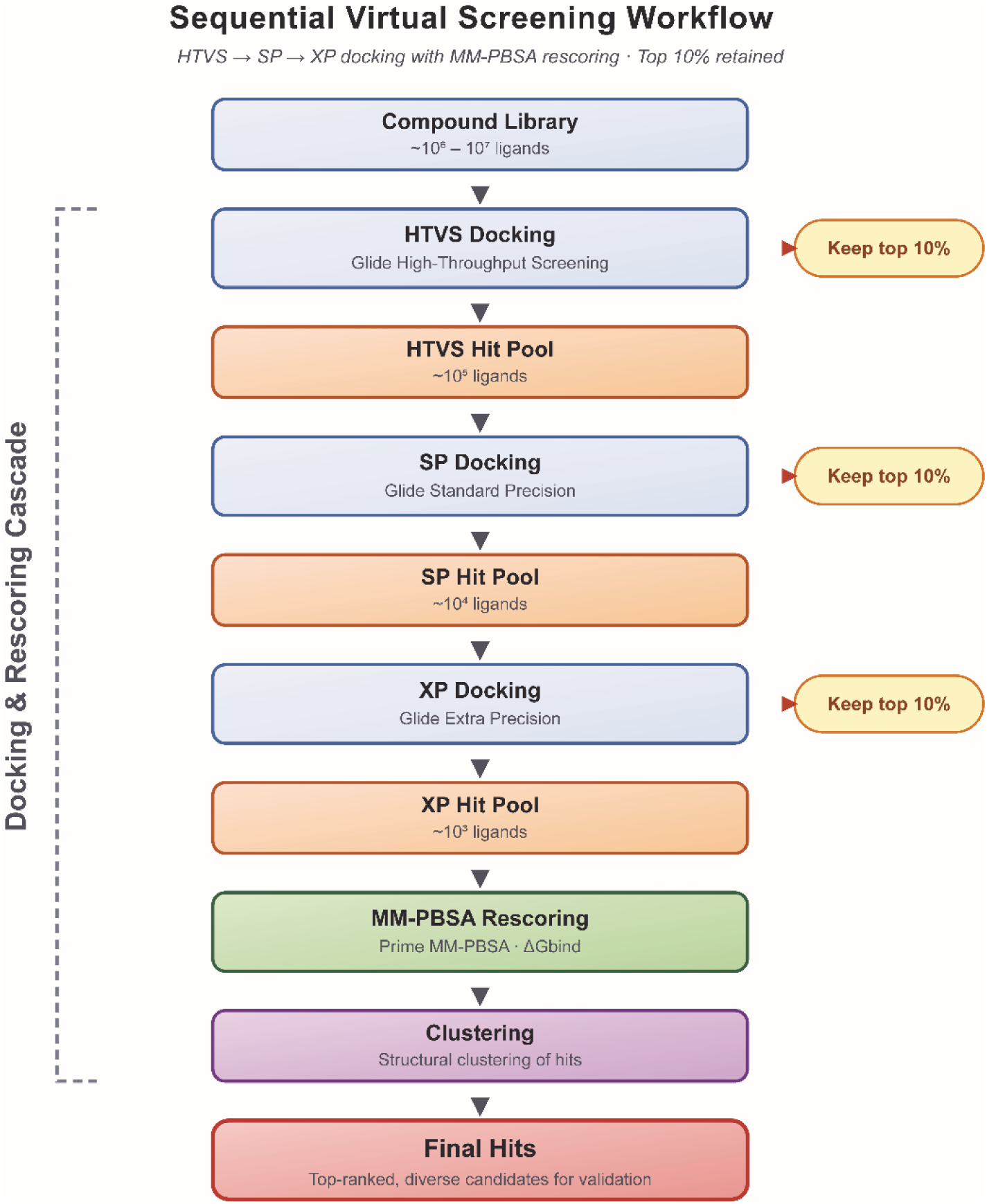
Virtual screening workflow of ANX B1 based drug discovery.

**Table 1.**
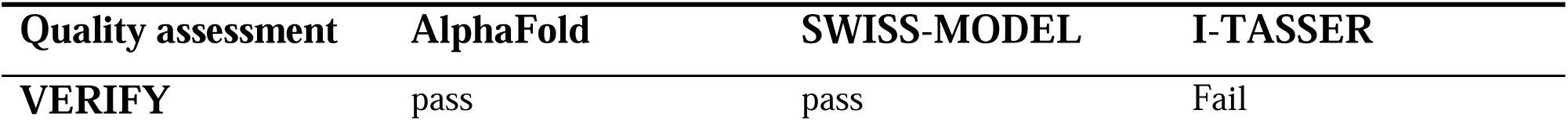
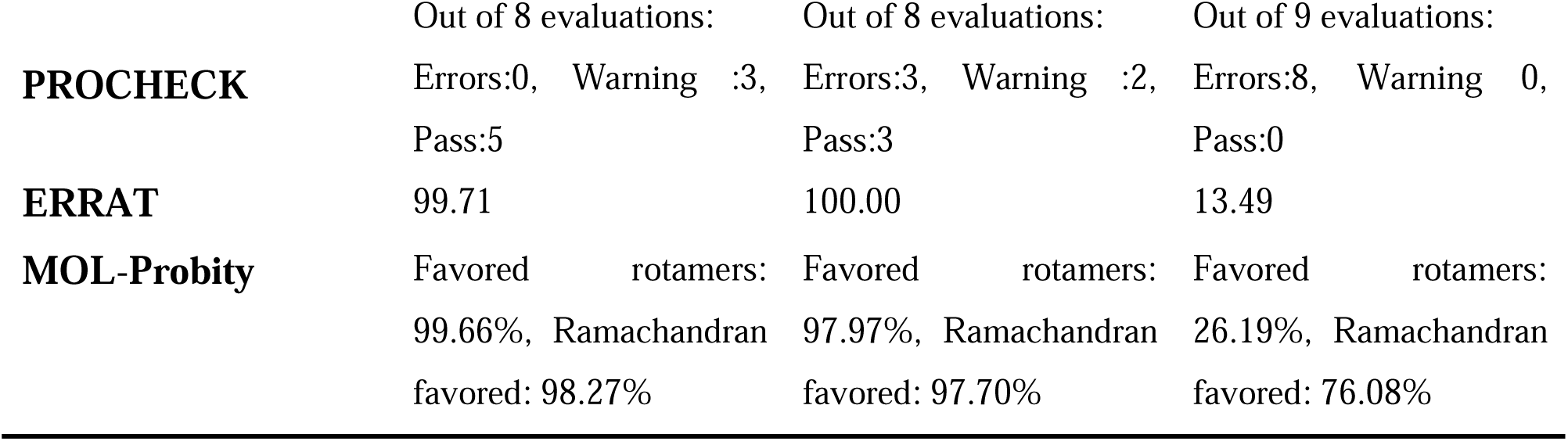
Protein model quality assessment based on SAVES and Mol-Probity.

A total of 1,456,161 input molecules were prepared for virtual drug screening using LigPrep, resulting in over 8,646,410 output molecules with Epik to add metal binding states. All output molecules were screened against ANX B1 through a sequential virtual screening workflow implemented in the Schrödinger suite, using HTVS, SP, and XP docking modes with MM/GBSA calculation (Figure 2). At each drug screen stage, only the top 10% of docked ligands were retained to progressively narrow the candidate pool toward high-affinity hits. The final XP docking step yielded 32 hit compounds, 11 commercially available compounds were selected for antiparasitic activity analysis as indicated in Table 1. These compounds displayed docking scores ranging from −11.174 kcal/mol to −7.526 kcal/mol, and their names, scoring values were summarized in Table 2.

**Table 2.** Compounds tested, anthelminthic efficacy, docking score, and maximum anthelmintic efficacy concentration.

| Compounds | Maximum<br>AE ( % ) | Concentration<br>causing the best AE<br>(mg/L) | XP_Docking<br>score (kcal/mol) | MM/GBSA<br>(kcal/mol) |
| --- | --- | --- | --- | --- |
| Abamectin | 100.00 | 0.2 | -9.701 | -43.91 |
| Amlodipine | 85.46 | 0.5 | -11.174 | -32.83 |
| Arecoline | 69.32 | 10 | -7.642 | -30.61 |
| Curcumin | 78.97 | 10 | -8.479 | -39.59 |
| Hydroxycitronellal | 28.75 | 10 | -8.519 | -31.28 |
| Pseudopelletierine | 58.25 | 10 | -8.103 | -33.27 |
| Pterostilbene | 58.33 | 10 | -7.955 | -33.35 |
| Polymyxin B Sulfate | 12.76 | 10 | -7.957 | -28.78 |
| Dihydrostreptomycin<br>Sulfate | 0 | - | -7.711 | -17.04 |
| Hydroxyethyl Cellulose | 0 | - | -7.526 | -16.1 |
| Imidazolidinyl Urea | 0 | - | -7.662 | -22.29 |

The *in vivo* anthelmintic performance of 11 compounds against *N. melleni* is presented in Table 2. Following 24 h of exposure, only abamectin with a docking score of −9.701 kcal/mol achieved complete parasite clearance, positioning it as the most effective candidates in this series. Although amlodipine preserved the lowest docking score (−11.174 kcal/mol), its anthelmintic efficacy was only 85.46% at higher concentration (0.5 mg/L) than that of abamectin. A similar trend was observed in the MM/GBSA values, where abamectin displayed a markedly more negative binding free energy (−43.91 kcal/mol) than amlodipine (−32.83 kcal/mol), consistent with its stronger *in vivo* activity. These findings indicate that, in this case, the MM/GBSA estimate aligned more closely with biological efficacy than the docking score alone, and they further highlight that individual computational parameters do not always correspond to *in vivo* performance. The observed mismatch between docking rankings and actual anthelmintic potency may reflect differences in target engagement, compound bioavailability, or off-target effects that are not fully captured by static scoring functions. No anthelmintic effect was detected in the DMSO-treated control group.

### *In vivo* effect of abamectin against *N. melleni*

The study evaluated the anthelmintic potential of abamectin against *N. melleni* and assessed its concurrent cytotoxicity profile. Morphological examination by optical microscopy confirmed the characteristic features of adult *N. melleni* individuals (Figure 4A), providing a baseline for subsequent drug response observations. Exposure to increasing concentrations of amlodipine produced a clear dose-dependent effect on parasite viability. Acute toxicity assays (Figure 4B) demonstrated that mortality rose progressively across the tested range of 0.01 to 0.50 mg/L, yielding a typical sigmoidal curve upon nonlinear regression fitting. Using a four-parameter DoseResp model (Figure 4C), the median lethal concentration (LC) was determined to be 0.254 mg/L (95% CI: 0.171-0.378 mg/L; R² = 0.940). This value establishes the upper boundary of safe drug exposure for the host organism under exposure conditions. Parallel anthelmintic efficacy tests (Figure 4D) revealed that amlodipine exerted potent antiparasitic activity at substantially lower concentrations. Efficacy increased steeply between 0.01 and 0.08 mg/L, with complete parasite elimination observed at concentrations ≥0.16 mg/L. The calculated median effective concentration (EC) was 0.033 mg/L (95% CI: 0.029-0.038 mg/L), indicating that half-maximal parasite clearance occurs at roughly one-eighth of the cytotoxic concentration. The disparity between these two values carries important implications for therapeutic application. The ratio of LC to EC yielded a therapeutic index (TI) of approximately 7.7, suggesting that amlodipine possesses a reasonable safety margin against *N. melleni* infection. A TI exceeding 5 is generally regarded as favorable in early-stage drug screening, as it implies that effective antiparasitic concentrations can be achieved without approaching host toxicity thresholds. From a mechanistic standpoint, the observed activity aligns with the established mode of action of macrocyclic lactones.[40] Abamectin acts primarily as a positive allosteric modulator of glutamate-gated channels[41, 42], which are widely expressed in invertebrate nervous systems. In platyhelminths, activation of these channels leads to sustained calcium ion influx, resulting in hyperpolarization of neuronal and muscle membranes.

**Figure 4.**
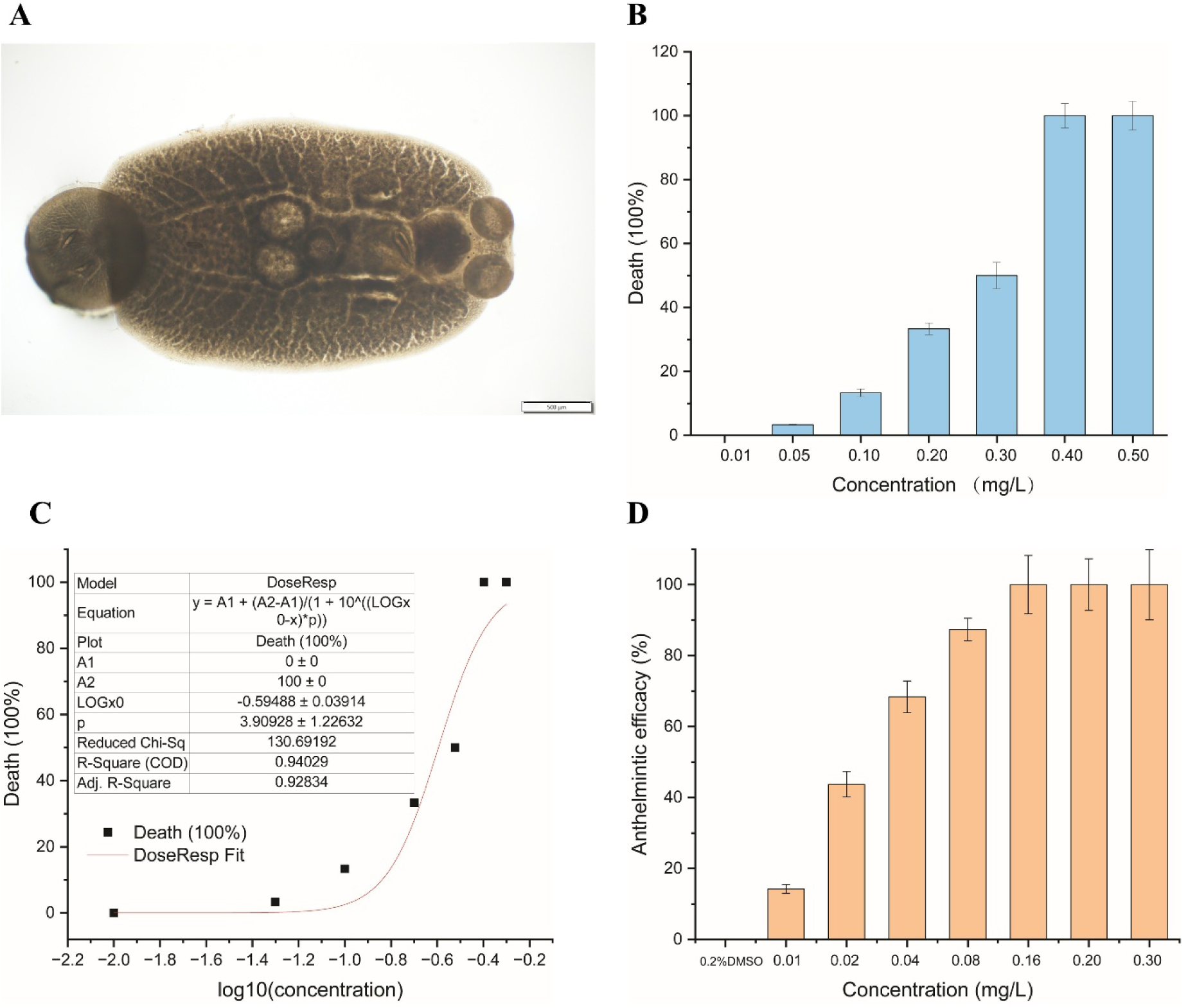
Optical microscope of *N. melleni* (A) and acute toxicity of abamectin to host fish (B), the nonlinear curve fitting to calculate LC_50_ (C) and its *in vivo* anthelmintic efficacy (D).

The present findings support abamectin as a potent antiparasitic agent against *N. melleni*. The EC value of 0.033 mg/L falls well below the toxic threshold, offering a practical safety margin for potential therapeutic use. Compared with conventional treatments for monogenean infections, this level of potency is competitive and warrants further investigation. Future studies should prioritize *in vivo* challenge trials using controlled infection models, accompanied by comprehensive histopathological evaluation of host organs. Pharmacokinetic profiling will be essential to determine whether waterborne or oral administration routes can maintain effective drug concentrations within the identified therapeutic window while avoiding accumulation to toxic levels.

### Molecular Dynamics simulation analysis of the annexin B1-abamectin complex over microsecond simulation

The molecular dynamics (MD) simulation of the annexin B1-abamectin complex was performed over 1000 ns to evaluate the structural stability, binding durability, and interaction mechanisms at the atomic level. As illustrated in Figure 5A, the root mean square deviation (RMSD) of the protein backbone (blue trace) exhibited an initial fluctuation, peaking around 150-200 ns, before stabilizing within the range of 1.8 to 2.4 Å for the remainder of the simulation. This plateau suggests that the protein backbone successfully relaxed into a stable conformation relative to the starting structure. Concurrently, the ligand RMSD (red trace) showed a sharp initial spike, likely corresponding to the induced-fit adjustment of abamectin within the binding pocket, before settling into a stable trajectory below 1.0 Å after 400 ns. The convergence of both traces confirms that the simulation reached equilibrium and that the complex is structurally robust. Figure 5B of RMSF analysis revealed distinct regions of mobility. Higher fluctuations were observed at the N-terminal region (residues 1-50) and specific loop regions (around residue 100 and 320), which is characteristic of the inherent flexibility of annexin family proteins. Conversely, the core helical regions remained relatively rigid, which is consistent with the secondary structure element (SSE) analysis in Figure 5D. The SSE plot demonstrates that the α-helical content remained constant at approximately 60% throughout the 1000 ns trajectory, with no signs of unfolding or denaturation, thereby validating the structural integrity of the protein during ligand binding.

**Figure 5.**
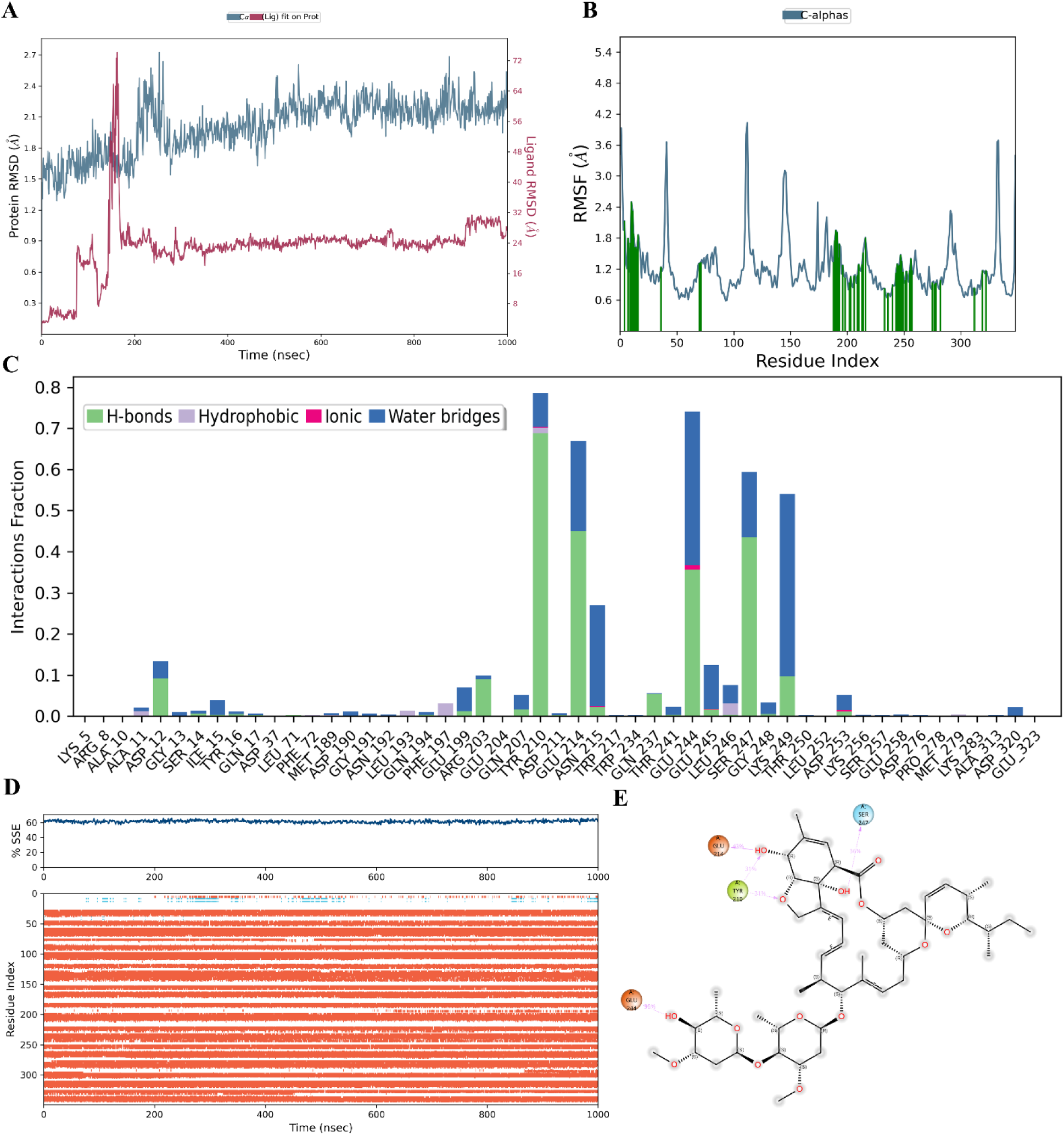
Molecular dynamics (MD) simulation and interaction analysis of the ANX B1-aba complex. **(A)** Time-evolution of the Protein RMSD (left Y-axis, blue line) and Ligand RMSD (right Y-axis, pink line) over a 1000-nsec simulation trajectory. **(B)** Root Mean Square Fluctuation (RMSF) of the C-alpha atoms as a function of the residue index, with significant interactive regions highlighted in green bars. **(C)** Categorized protein-ligand interactions monitored throughout the simulation, displaying the interaction fraction of specific residues participating in hydrogen bonds (green), hydrophobic contacts (grey), ionic interactions (pink), and water bridges (blue). **(D)** Secondary structure element (SSE) composition and stability analysis. The top panel shows the total percentage of SSE (%SSE) over time, and the bottom panel tracks the evolution of specific secondary structure assignments for each amino acid residue across the 1000-nsec trajectory. **(E)** Detailed 2D schematic representation of the specific molecular interactions and contact network between the ligand and key binding site residues of the target protein.

The interaction profile between annexin B1 and abamectin was dominated by hydrophobic contacts and water bridges, as quantified in Figure 5C. While hydrogen bonds (green bars) contributed to specificity, involving residues such as TYR-210, GLU-214, GLU-244, and SER-247. Notably, water bridges (dark blue) played a significant role in mediating interactions at residues like ASN-215, GLU-244, and LYS-249, suggesting that the binding pocket utilizes structural water molecules to enhance complementarity. The contact frequency map (consistent with previous contact analysis) indicates that once the ligand settled (post-200 ns), these interactions were maintained with high occupancy. The 3D interaction diagram (Figure 5E) provides a visual confirmation of the binding mode. Abamectin’s macrolide ring system is deeply embedded in a hydrophobic cleft formed by the annexin B1 helix bundle. Key polar contacts were identified with residues such as TYR-210, GLU-214, and SER-247, which may stabilize the lactone moiety. The presence of GLU-244 near the hydrophobic tail of abamectin further locks the ligand in place. The stability of these interactions over the microsecond timescale supports the hypothesis that annexin B1 is a plausible drug target for abamectin. Unlike typical ion channel targets, this interaction suggests a potential modulatory role on annexin B1’s function-perhaps interfering with its membrane aggregation or calcium channel properties-which could contribute to the anthelmintic efficacy observed in *N. melleni*.

### Conclusion

In this study, we established a structure-based virtual screening strategy centered on *Neobenedenia melleni* annexin B1 for the discovery of anti-monogenean compounds in pearl grouper aquaculture. Phylogenetic analysis confirmed that annexin B1 is a parasite-specific protein clearly distinct from host grouper annexins, justifying its use as a molecular target. A hierarchical in silico pipeline-combining AlphaFold-based structure prediction, docking of a 1.45-million-compound library (HTVS/SP/XP), MM/GBSA rescoring, and ADMET filtering-yielded eleven candidates for *in vivo* validation. Among these, abamectin exhibited the most potent anthelmintic activity, achieving complete parasite clearance with an EC_50_ of 0.033 mg/L and a therapeutic index of approximately 7.7 (host 24-h LC_50_ = 0.254 mg/L). Molecular dynamics simulations (1000 ns) and MM/GBSA analysis demonstrated that abamectin binds annexin B1 through stable hydrophobic and water-bridged polar interactions, supporting the protein as a bona fide target and rationalizing its antiparasitic efficacy. Together, these findings establish a generalizable, target-directed framework for antiparasitic drug discovery in aquaculture and identify abamectin as a promising, low-toxicity lead compound. Future work should prioritize *in vivo* challenge trials under controlled infection, pharmacokinetic profiling to optimize administration route and dose, and evaluation of field-scale safety and environmental fate.

## ASSOCIATED CONTENT

### CRediT authorship contribution statement

Longkun Gao, Wei Luo: Writing – original draft, Visualization, Methodology, Investigation. Yanru Guo: Methodology, Formal analysis. Ying Yan, Guanhai Li: Methodology, Visualization. Qin Yu, Minzhu Liu: Formal analysis. Erlong Wang: Formal analysis, Conceptualization. Pengfei Li: Resources, Conceptualization. Tianqiang Liu: Project administration, Funding acquisition, Writing – review & editing

## Acknowledgements

We thank Guangxi Science and Technology Program (Grant no. 2023GXNSFAA026500), Shenzhen Science and Technology Program (Grant no. JCYJ20240813151900002), and Key Research and Development Program of Shaanxi (Program no. 2024PT-ZCK-02) for supporting this work. We also thank the help of Miss Min Zhou from the Northwest A&F University’s Life Science Core Services facility with the assistance in obtaining relevant images.

## Declaration on the Use of AI in Scientific Writing

The authors developed the core concepts, data, and initial drafts of this manuscript. DeepSeek V4 Pro (https://www.deepseek.com/) was subsequently employed to optimize the text flow, reorganize paragraph structures, and enhance grammatical accuracy. The final version was carefully verified, revised, and approved by all authors, who accept sole responsibility for the integrity of the work.

